# A knock-in *Drosophila* model supports a conserved link between potassium channelopathy and involuntary movement

**DOI:** 10.1101/2020.02.20.957571

**Authors:** Patrick Kratschmer, Edgar Buhl, Ko-Fan Chen, Simon Lowe, Dimitri M. Kullmann, James J.L. Hodge, James E.C. Jepson

**Affiliations:** Department of Clinical and Experimental Epilepsy, UCL Queen Square Institute of Neurology, London, UK; School of Physiology, Pharmacology and Neuroscience, University of Bristol, Bristol, UK

**Keywords:** *Drosophila*, BK channel, Slowpoke, paroxysmal dyskinesia

## Abstract

**Background:** Genetic and in vitro studies have linked a heterozygous gain-of-function mutation (D434G) in the hSlo1 BK (Big potassium) channel to paroxysmal dyskinesia. However, support for this linkage from in vivo models has been lacking.

**Objectives:** We aimed to re-create the equivalent mutation to hSlo1 D434G in the fruit fly, *Drosophila*, and examine how this mutation altered movement and action potential waveforms.

**Methods:** We generated a knock-in *Drosophila* model of hSlo1 D434G. We used video-tracking and infra-red beam-break systems to test whether locomotion was altered in this model, and patch-clamp electrophysiology to determine how the mutation affected action potential waveforms.

**Results:** We identified profound motor dysfunction and sporadic leg twitches, as well as a reduced width and an enhancement of the afterhyperpolarization phase of action potentials, in the model background.

**Conclusion:** Our results support a conserved relationship between enhanced BK channel function and disrupted motor control across distantly related species.

## Introduction

Paroxysmal dyskinesias (PxDs) are characterised by intermittent involuntary dystonic, choreiform and/or ballistic movements^1^. A variety of mutations have been linked to inherited PxDs, providing a platform to uncover cellular and circuit-level alterations underlying these disorders^1-3^. One form of PxD with a compelling molecular causation is *KCNMA1*-linked paroxysmal non-kinesigenic dyskinesia, termed PNKD3 (OMIM# 609446)^4 5^. Clinical features of PNKD3 include dystonic and choreiform movements of the mouth, tongue and extremities that can occur spontaneously but are also triggered by fatigue, stress and alcohol^5^. In some patients dyskinesia is co-morbid with absence or generalised tonic-clonic seizures^5^.

Prior work identified an autosomal dominant mutation (1301A→G) in the *KCNMA1* locus that co-segregated with PNKD3 in a multi-generation family^5^. *KCNMA1* encodes the α-subunit of the BK potassium channel (hSlo1). BK channels are activated by membrane depolarisation and increased intracellular Ca^2+^, and play conserved roles in modulating the repolarisation and fast afterhyperpolarization (AHP) phases of action potentials (APs) in both vertebrate and invertebrate neurons^6-9^. The 1301A→G mutation replaces a negatively charged aspartic acid (D) at residue 434 with a neutral glycine (G)^5^, and in vitro analyses in non-excitable cells suggest that D434G acts as a gain-of-function (GOF) mutation by enhancing Ca^2+^-sensitivity, increasing the rate of channel activation, and slowing deactivation^5 10 11^.

While distinct mutations in *KCNMA1* have been linked to PxD with partially overlapping clinical characteristics^12^, the D434G mutation has yet to be independently identified in other pedigrees. Thus, support from in vivo models linking equivalents of this mutation to altered movement would strengthen the connection between genotype and pathological phenotype in PNKD3. In addition, how D434G impacts the dynamics of APs in neurons has yet to be explored. Here we utilise *Drosophila* to test whether the equivalent mutation to hSlo1 D434G disrupts motor control and alters AP waveforms in a manner consistent with BK channel GOF.

## Materials and Methods

### *Drosophila* husbandry

Flies were maintained on standard fly food at constant temperature 25°C under 12 h: 12 h light-dark cycles (12L: 12D). The following strains were obtained from the Bloomington stock center: *y, w; hs-FLP, hs-I-SceI*/CyO (BDSC #6934), *y, w, ey-FLP* (BDSC #5580) and *y, w, Cre; +; D*/*TM3, *sb* (BDSC #851). The isogenic iso31 wild type strain used for outcrossing and *dysc*^s168^ mutants were kind gifts from Kyunghee Koh (Thomas Jefferson University). *PDF∷RFP* was a kind gift from Dr Justin Blau (New York University), See Supplemental Information for further details of Materials and Methods.

## Results

The D434 residue mutated in PNKD3 is located within the regulator of K^+^ conductance 1 (RCK1) domain of the channel^5^ (Fig. 1A), which contains binding sites for divalent cations and functions to connect Ca^2+^-binding to channel opening^11^. Consistent with its functional importance^5 10 11^, the D434 residue is highly conserved across Bilateria^13^ (Fig. 1B). D appears fixed in Deuterostomes at equivalent positions to hSlo1 434, while in Protostome orthologs, including the *Drosophila* BK channel α-subunit Slowpoke (SLO), a glutamic acid residue (E) is more prevalent (Fig. 1B). Importantly, mutating the murine equivalent of D434 (D369 in mSlo1) to E does not alter channel function at physiological Ca^2+^ concentrations^11^, consistent with the similar physiochemical properties of aspartic and glutamic acids. Since the above evidence supports functional conservation of this residue between humans and *Drosophila*, we used ends-out homologous recombination to substitute the *Drosophila* residue orthologous to hSlo1 D434 (SLO E366) with glycine (Supplemental Figs. 1 and 2), generating an invertebrate model of PNKD3. In parallel we isolated corresponding controls harbouring the genomically encoded E residue (Supplemental Figs. 1 and 2). As part of the homologous recombination process, both lines contain a 76 bp sequence in a non-conserved intronic region of *slo* that includes a single *loxP* site^14^ (Supplemental Fig. 1). We isolated ten E366G and four control alleles, and subsequently out-crossed three of each to an isogenic iso31 strain for five generations to homogenize genetic background. Since D434G is dominant we analysed heterozygotes for the E366G and control alleles. We term these flies *slo*^E366G/+^ and *slo*^*loxP*/+^ respectively.

**FIG 1.**
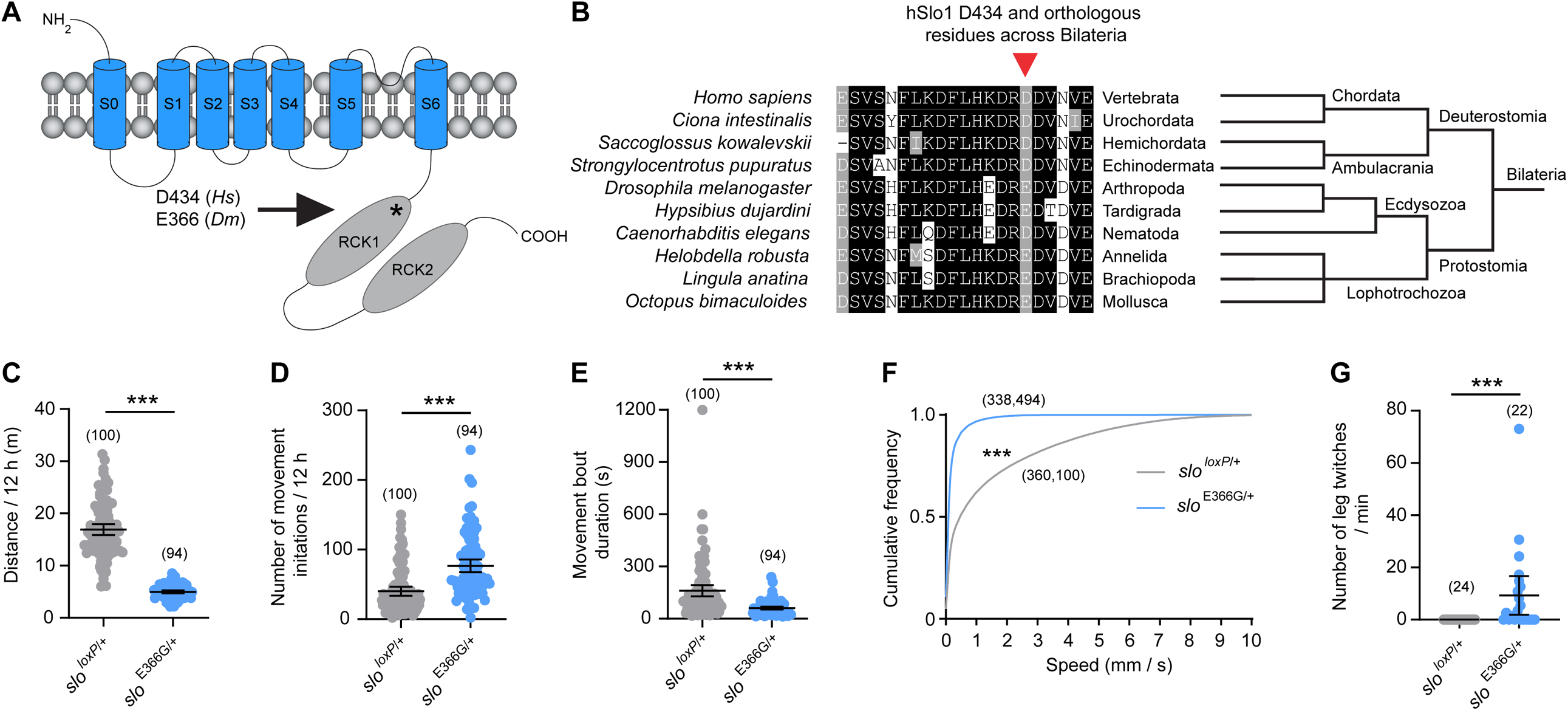
(A) Schematic showing approximate location of the D434 and E366 residues in the RCK1 domain of human and *Drosophila* hSlo1/SLO. (B) Alignment of residues surrounding hSlo1 D434 (arrow) with orthologous BK α-subunits spanning > 540 MY of evolutionary divergence^13^. (C-F) Movement parameters in adult male *slo*^*loxP*/+^ and *slo*^E366G/+^ flies measured by the DART system. (G) Frequency of leg twitches in adult male *slo*^*loxP*/+^ and *slo*^E366G/+^ flies. Dots in panels C, D, E, and G represent individual flies; n-values are noted. Error bars: 95% Confidence Interval (CI). ***p<0.0005, Mann-Whitney U-test (C, D, E, G) or Kolmogorov-Smirnov test (F).

**FIG 2.**
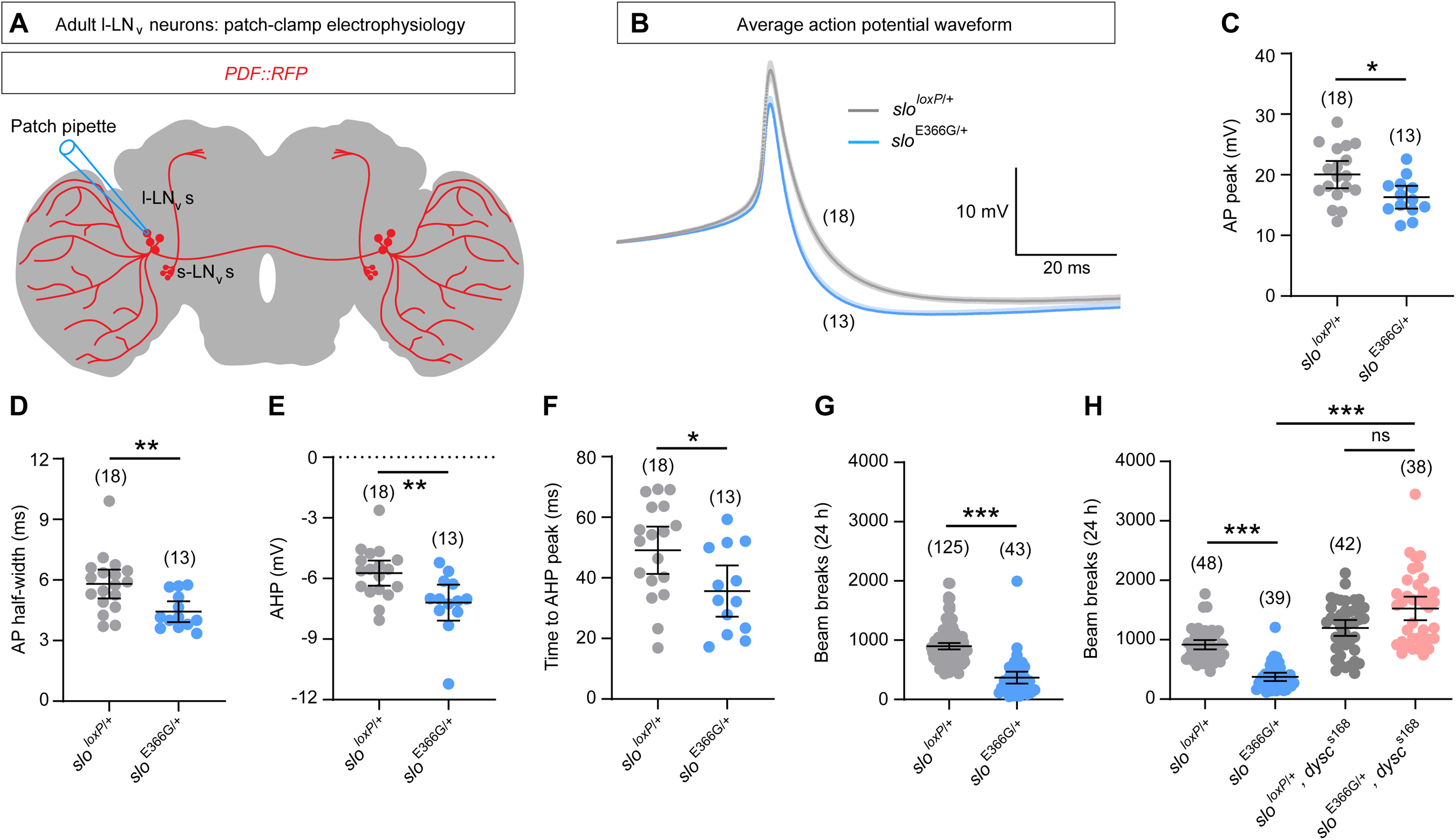
(A) Illustration showing morphology of l-LN_v_s^26^ labelled with *PDF* promoter-driven RFP (*PDF∷RFP*^*27*^) and location of patch-clamp recording sites. (B) Average action potential (AP) waveforms in l-LN_v_s. Darker and lighter shades show mean and standard error of the mean (SEM). (C-F) l-LN_v_ AP and AHP parameters. (G-H) Total activity recorded by the DAM system over 24 h. n-values are noted. Error bars: 95% CI. *p<0.05, **p<0.005, ***p<0.0005, ns – p>0.05, unpaired t-test with Welch’s correction (C, D, F), Mann-Whitney U-test (E, G), or Kruskal-Wallis test with Dunn’s post-hoc test (H).

We addressed two key questions relating to the role of BK channels in PNKD3. Firstly, does heterozygosity for the *slo*^E366G^ allele alter movement in *Drosophila*? Secondly, is the impact of this mutation on neuronal physiology and behavior consistent with a GOF effect? To address the first question, we used an automated video-tracking system (*Drosophila* Arousal Tracking; DART^15^) to monitor locomotion in adult male *slo*^E366G/+^ and *slo*^*loxP*/+^ flies. *slo*^E366G/+^ adult males exhibited a robust reduction in distance travelled over a 12 h period compared to controls (Fig. 1C), an effect consistent across independently derived recombinant lines (Supplemental Fig. 3). Since male flies are inactive for much of the day^16^, we performed a detailed analysis of movement during a period of normally heightened activity: 0-1 h following lights-on in 12 h light: 12 h dark conditions. During this time span, we found that *slo*^E366G/+^ males initiate movement more frequently than controls (Fig. 1D). However, the duration of locomotor bouts was shorter, and overall locomotor speeds were significantly reduced, in *slo*^E366G/+^ males (Fig. 1E, F). Similar results were observed in *slo*^E366G/+^ females (Supplemental Fig. 4). Strikingly, upon detailed analysis of videos we also observed that many *slo*^E366G/+^ flies exhibited bouts of spontaneous leg-twitches (median duration: 3 s) reminiscent of jerk-like dyskinetic movements, and which were absent in controls (Fig. 1G, Supplemental Fig. 5 and Supplemental Video 1).

Since motor control was clearly perturbed in *slo*^E366G/+^ flies, we next assessed how this mutation impacted neuronal physiology. We performed ex vivo patch-clamp recordings from a cluster of SLO-expressing neurons termed large ventral lateral neurons (l-LN_v_s)^17^, which can be identified based on their expression of the neuropeptide Pigment Dispersing Factor (PDF) (Fig. 2A). Passive membrane properties of l-LN_v_s were not altered in *slo*^E366G/+^ flies (Supplemental Fig. 6A, B). In contrast, analysis of spontaneous APs demonstrated a significant reduction in mean AP amplitude and duration, as well as enhanced AHP amplitude and accelerated AHP kinetics, in the *slo*^E366G/+^ background (Fig. 2B-F). These findings are consistent with an enhancement of BK channel function in *slo*^E366G/+^ neurons^7-9^. Interestingly, while the D434G mutation has been hypothesized to increase neuronal firing rates by enhancing the rate of sodium channel recovery^5^, we did not observe an alteration in either the rate of spontaneous firing (Supplemental Fig. 6C) or of higher frequency firing induced by +20 or +40 pA current injections (Supplemental Fig. 7) in *slo*^E366G/+^ l-LN_v_s.

Finally, we sought to provide evidence that the E366G mutation causes SLO GOF at the organismal level. We recently identified a regulator of SLO called Dyschronic (DYSC), a PDZ-domain-containing ortholog of the human protein Whirlin that is mutated in the deaf-blindness disorder Usher syndrome^18 19^. DYSC binds to and promotes SLO expression in *Drosophila* neurons, and *dysc* mutants exhibit drastically reduced neuronal SLO levels and associated currents^18^. Loss of DYSC disrupts clock-driven alterations in locomotion in adult *Drosophila*, but gross motor capacity appears largely unimpaired in *dysc* mutants^18^. We therefore hypothesized that loss of DYSC might suppress motor dysfunction in *slo*^E366G/+^ flies by reducing the expression of pathogenic GOF SLO. To test this we utilised a distinct locomotor analysis platform, the *Drosophila* Activity Monitor (DAM), which enables automated recordings at higher throughput relative to DART^20^. We confirmed that reduced locomotion in *slo*^E366G/+^ flies can be quantified using the DAM (Fig. 2G), then examined whether reduced DYSC levels (via homozygosity for the loss-of-function allele *dysc*^s168 18^) could rescue this phenotype. Indeed, we observed a restoration of locomotor activity in *slo*^E366G/+^, *dysc*^s168^ double-mutants (Fig. 2H). Collectively, the above data provide evidence at both the cellular and organismal levels that the *Drosophila* equivalent of hSlo1 D434G enhances BK channel function in vivo and impairs movement control.

## Discussion

Here we present a *Drosophila* model of PNKD3 dyskinesia. The multifaceted alterations in motor control and sporadic leg twitches in *slo*^E366G/+^ flies support the genetic linkage of the *KCNMA1* D434G mutation with dyskinesia^5^ and suggest that BK potassium channels play conserved roles in the regulation of movement across Bilateria.

We distinguish two movement-related effects of the *slo*^E366G/+^ mutation: a highly penetrant perturbation of overall locomotor capacity, and a more variable phenotype consisting of sporadic bouts of leg twitches. In contrast, gross locomotor ability in PNKD3 patients is largely normal, with infrequent dyskinetic attacks generally triggered by alcohol, stress or fatigue^5^. Such differences in phenotypic penetrance and severity between *Drosophila* and humans may arise through several interlinking mechanisms, including the absence of modulatory BK channel β-subunits in *Drosophila* (known to alter the impact of the PNKD3 mutation on mammalian BK channel function^10 21^), and species-specific differences in the expression of other ion channels contributing to AP dynamics as well as overall neural architecture. Nonetheless, the robust motor defects observed in *slo*^E366G/+^ flies are an advantageous aspect of this model, since high-throughput automated systems such as the DAM can now be deployed in concert with classical genetics to search for conserved genetic modifiers of these phenotypes.

The SLO E366G mutation alters neuronal AP waveforms in a manner consistent with BK channel GOF^7^, supporting an array of detailed studies examining how the D434G mutation and murine equivalents influence BK channel properties in non-neuronal cells^5 10 11 21 22^. However, further investigations are required to examine how these changes in AP dynamics impact overall synaptic output. Mammalian BK channels can both inhibit or enhance neurotransmitter release depending on cell-type^4 23-25^. Thus, elucidating the patho-mechanisms underlying PNKD3 requires an understanding of both the neural subpopulations driving movement defects and how BK channel GOF alters their excitability. Our work validates *slo*^E366G/+^ flies as a model to undertake such studies. Indeed, in our accompanying paper we exploit unique aspects of the larval stage of the *Drosophila* lifecycle to demonstrate that the *slo*^E366G/+^ mutation can reduce neurotransmitter release, alter short-term synaptic plasticity, and disrupt the pre-motor central pattern generator that drives larval foraging behavior. More broadly, our results demonstrate that *Drosophila* can be utilised as a platform to confirm genotype-phenotype correlations in PxDs and interrogate the neural basis of these disorders.

## Acknowledgements

We thank Prof. Kyunghee Koh (Thomas Jefferson University) and Prof. Justin Blau (New York University) for sharing *Drosophila* stocks; Prof. Henry Houlden and Talya Goble (University College London) for comments on the manuscript; and Tom Lowe for help with video editing.

## Author Contributions

Conceptualization: P.K., J.E.C.J. Methodology: P.K., E.B., K-F.C., S.L., J.E.C.J. Software: P.K. Validation: P.K., E.B. Formal Analysis: P.K., E.B., S.L., J.E.C.J. Investigation: P.K., E.B., K-F.C., S.L. Writing – Original Draft: J.E.C.J. Writing – Review and Editing: P.K, E.B, K-F.C, S.L, D.M.K, J.J.L.H., J.E.C.J. Visualisation: P.K., E.B., S.L., and J.E.C.J. Supervision: D.M.K, J.J.L.H., J.E.C.J. Project Administration: J.J.L.H., J.E.C.J. Funding Acquisition: P.K., D.M.K, J.J.L.H., J.E.C.J.

## Relevant conflicts of interest/financial disclosures

None for all authors.

## Supplemental Information, Figures and Video Legends

### Supplemental Materials and Methods

#### Generation of the *slo*^E366G^ and *slo*^*loxP*^ alleles

Ends-out homologous recombination was performed as described previously^1 2^. Recombinogenic arms corresponding to the *slo* locus were amplified using the following primers: Arm 1 forward – <u>CGTACG</u>TCCCCAAGTACAGACAGCAA, Arm 1 reverse – <u>GGCGCGCC</u>GTTGTCAGTGTGTCGTGTGC; Arm 2 forward – <u>GGTACC</u>GCAGCTCAATGGAATGTGATT, Arm 2 reverse – <u>GCGGCCGC</u>ACGCTTATTCTGGGACTTCG. Underlined sequences indicate restriction enzyme cut sites^2^. The above primers amplify 2491 bp 5’ of position chr3R:24,686,818 (*Drosophila melanogaster* Genome Assembly BDGP6) for Arm 1 and 2731 bp 3’ of position chr3R:24,686,862 for Arm 2. The A1097G point mutation resulting in the E366G amino-acid change was introduced into Arm 2 via a customised DNA fragment (GeneArt Gene Synthesis, ThermoFisher Scientific) that was used to replace the 5’ 722 bp of Arm 2. Arms were cloned into *p[w25*.*2]* using standard techniques. Embryonic injection of the *p[w25*.*2]*-Arm2^E366G^-Arm1 vector was performed by BestGene Inc. (CA, USA). Primers used to validate successful recombination events (Supplemental Fig. 1B) are as follows: pW-Acc3_F (forward primer) – GCTCAGCTTGCTTCGCGATGTGTTCAC, pW-Acc1_R (reverse primer) – TTAGTTGAGTGCTTAAATTCAAAGGAT. The presence or absence of the A1097G point mutation in Arm 2 was verified via Sanger sequencing using the following primers: a2_alternative_R2-validation_F (R2F) – TCCGCTTTAATCGCACACTA; GEPD_seq_1 – CCCCCACCTTCAACAACACA. We also sequenced the entirety of Arm 1 and Arm 2 post-recombination and did not find any additional sequence variants in these regions apart from previously described intronic polymorphisms^3 4^. To outcross each allele into an isogenic background, the following primers were used: OCF (forward primer) *–* AGACTAGTCTAGGGTACCGCA, OCR (reverse primer) – TAGTTCCTTGAATTGGCAGCG. OCF binds to a portion of the 76 bp of sequence incorporated into the intron upstream of exon 10 (which includes the single *loxP* site), and therefore will only generate a product in a *slo*^E366G/+^ or *slo*^*loxP*/+^ background.

#### *Drosophila* behavioural analyses

For behavioural analyses, three- to seven-day old male or female flies were collected and loaded into glass tubes (Trikinetics inc., MA, USA) containing 4% sucrose and 2% agar (w/v). Locomotor activity was recorded using the *Drosophila* Activity Monitor (DAM, TriKinetics inc. MA, USA) system or *Drosophila* ARousal Tracking (DART, BFKlab, UK) in 12L: 12D at 25^°^C as described previously^5^. A customised R script (R v. 3.6.0) was used to analyse movement parameters derived from the DART video-tracking data, available on GitHub: https://github.com/PatrickKratsch/. This script first creates an ‘offset table’ from the DART tracking data, obtained by subtracting successive x- and y-values. Hence, the offset table defines the displacement for each fly between consecutive frames. The total distance travelled between consecutive frames was calculated for each fly. Summing all frame-to-frame displacements over 12 h yielded the total distance travelled during this period. Flies that did not move during the second half of the experiment (hours 7 to 12) were identified as dead and excluded from further analysis. Of note, the precise duration analysed was 716 min (11 h and 56 min) due to the removal of 2 min post-lights-on and 2 min pre-lights-off, during which the lighting conditions prevented appropriate video-tracking. Analyses of other movement parameters are based on a period of 1 h after the dark-light transition (9:02 am – 10:02 am), a time of robust activity in wild-type flies. To analyse the distribution of locomotor speed, empirical cumulative distribution plots were generated based on 1 s movement bins (3601 bins for a 1 h and 1 s period). To analyse movement initiation, a sliding window of length 2 quantified how often a 1 s bin of non-movement was followed by a 1 s bin of movement. To analyse mean movement bout length, the fly movement data were transformed into a binary matrix, with 1’s indicating movement and 0’s non-movement. Subsequently, the lengths of movement streaks (streaks of 1’s) were extracted and averaged. DAM data were analysed using a customised R script, which is available on GitHub: https://github.com/PatrickKratsch/. This script reads the DAM output (TXT files) for each monitor into separate data objects and extracts the relevant experimental time. It then extracts the beam break data for each fly and sums the corresponding values over 24 h, yielding the total number of beam breaks per fly. Flies that did not move at all (0 beam breaks) during the second 12 h of the experiment were identified as dead and removed from further analyses. To quantify spontaneous leg-twitches, manually recorded videos were taken using an iPad Pro (Apple, CA, USA). Glass tubes (Trikinetics) were placed horizontally to avoid any potential confounding effects of possible alterations in geotaxis. Videos were analysed blind with respect to genotype. To ensure that our analysis did not conflate ‘dyskinetic movements’ with other motor behaviors such as grooming or locomotion, leg twitches were scored using the following criteria. 1. That leg ‘twitches’ occurred only in a single limb; instances where >1 limb exhibited simultaneous movement were not scored. 2. That each bout consisted of >1 repetitive movements of similar characteristics. 3. That leg movements did not involve grooming of any body part. 4. That movements were not coincident with forward or backwards locomotion of > 1/2 body length. 5. Where a fly was positioned on the agar/sucrose food or attached to the cotton wool at the opposite length of the tube, movements were also not scored. When a bout of leg twitches occurred, videos were assessed in a frame-by-frame manner (frame rate: 30 Hz) to quantify the number of twitches during the bout.

### Patch-clamp electrophysiology

Whole-cell patch clamp recordings were performed on large lateral ventral neurons (l-LN_v_) as described previously^6^. To visualize the neurons a *PDF∷RFP* fusion construct^7^ was used and crossed to *slo*^*loxP/loxP*^ homozygotes for control and *slo*^E366G^*/TM6b, tb* for the experimental genotype. Two- to five-day old male flies were decapitated at ZT18-20 (i.e. 6-8 h after lights-off) under red light illumination and brains dissected in external solution (in mM: 101 NaCl, 1 CaCl_2_, 4 MgCl_2_, 3 KCl, 5 glucose, 1.25 NaH_2_PO_4_, 20.7 NaHCO_3_, pH 7.2). This time-point has previously been shown to represent a period of maximal SLO expression in the l-LN_v_s^8^. After cleaning and removal of the ganglion sheath, brains were placed in the recording chamber ventral side up and secured using a custom-made harp. Recordings were made at room temperature (20-22°C) with borosilicate glass electrodes (8-15 MΩ resistance) filled with internal solution (in mM: 102 K-gluconate, 17 NaCl, 0.94 EGTA, 8.5 HEPES, 0.085 CaCl_2_, 1.7 MgCl_2_, pH 7.2). Signals were amplified (Axon MultiClamp 700B), digitized (Axon DigiData 1440A; sampling rate: 20 kHz; filter: Bessel 10 kHz), recorded (pClamp 10, Molecular Devices, Sunnyvale, CA, USA) and the liquid junction potential (13 mV) subtracted from the membrane voltages before analysis. The resting membrane potential (RMP) and spontaneous firing rate (SFR) were measured after allowing the recordings to stabilize for one minute. Input resistance was calculated using Ohm’s law by measuring the voltage change in response to hyperpolarizing currents and excitability was measured by injecting depolarizing current pulses (0-40 pA, 5 pA increments) of either 1 s or 5 s duration. The action potential size (AP peak), width at half maximal amplitude (half-width), afterhyperpolarization (AHP) amplitude and the time from peak to AHP were measured from averages of 10 spikes for each recording, aligned to the peak and measured relative to RMP.

### Bioinformatics

hSlo1 orthologs were initially identified using BLASTp (https://blast.ncbi.nlm.nih.gov/Blast.cgi?PAGE=Proteins), using either the hSlo1 21 amino-acid sequence shown in Fig. 1A or full-length *Drosophila* SLO as queries. Orthologous hits defined as encoding Ca^2+^-activated K^+^ channels were aligned in Clustal Omega (https://www.ebi.ac.uk/Tools/msa/clustalo/). Alignments were visualised using BoxShade (https://embnet.vital-it.ch/software/BOX_form.html), with 70% of sequences required to agree for shading. Black shading indicates amino-acid sequence conservation; grey shading indicates functional conservation. Bilateral phylogenetic tree (Fig. 1B) is derived from Telford et al., (2015)^9^. Selected clades within the Protostome lineage are shown.

### Statistics

Data populations were initially examined for normal distributions using the Shapiro-Wilk test. Statistical differences in normally distributed populations were tested for via unpaired t-tests with Welch’s correction for non-identical variance. Non-normal populations were assessed via Mann-Whitney U-test or Kruskal-Wallis test with Dunn’s post-hoc test. Quantification of leg-twitches in Fig. 1G were analysed blind with respect to genotype. For locomotor analyses in Figs. 1C-F we pooled individuals from three independent recombinant *slo*^E366G/+^ and *slo*^*loxP*/+^ lines (Supplemental Fig. 2). Since individual recombinant lines did not show any differences in overall movement (Supplemental Fig. 3), video analysis (Fig. 1G), patch-clamp electrophysiology (Fig. 2B-F) and DAM studies (Fig. 2G, H) were compared between flies from single representative *slo*^E366G/+^ and *slo*^*loxP*/+^ lines (25.1.1 and 132.1.1; Supplemental Figs. 2 and 3). All DAM and DART datasets were derived from ≥ 3 independent biological replicates.

## Supplemental Figures and Figure Legends

**Supplemental Fig. 1.**
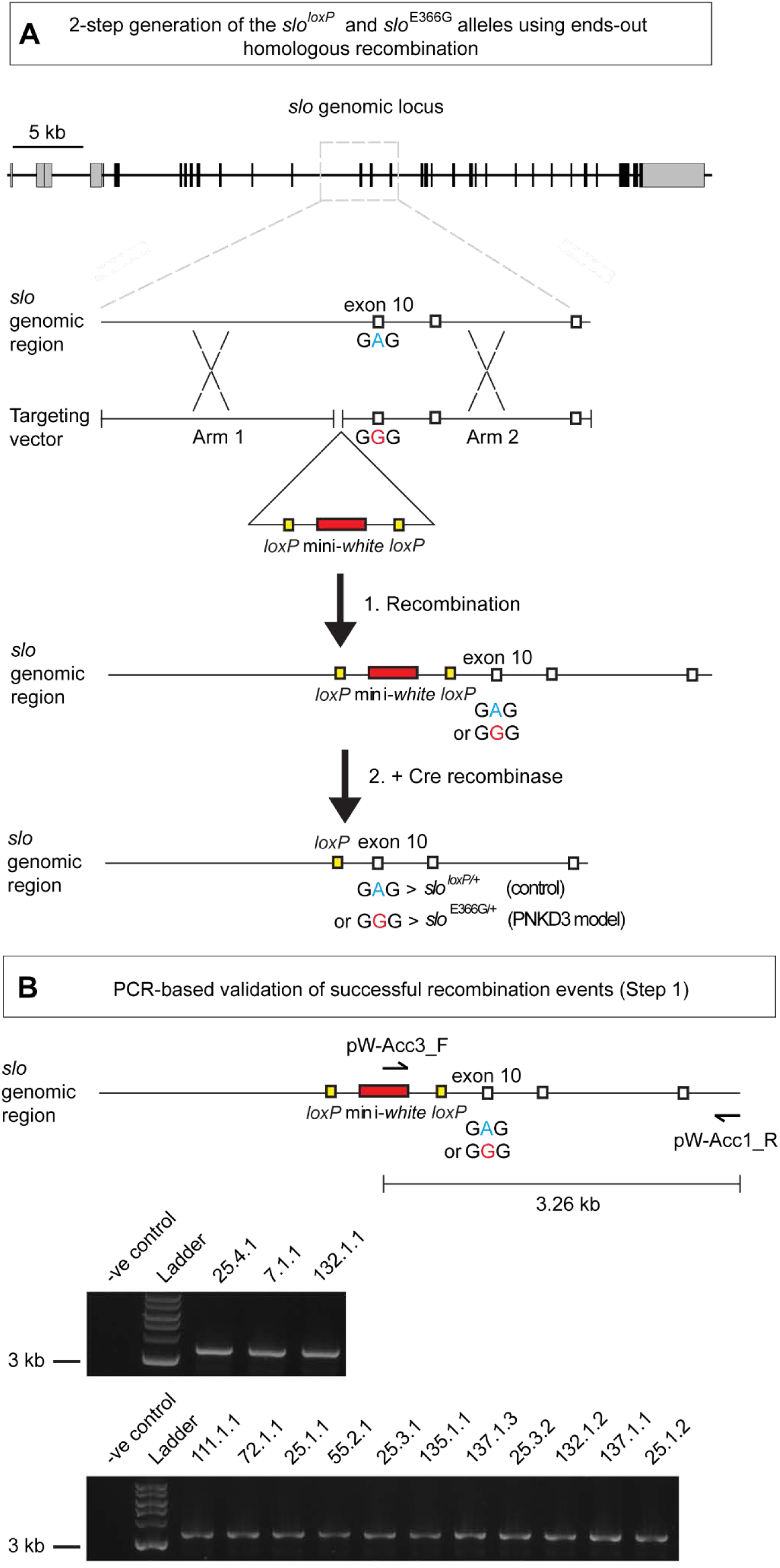
(A-B) Schematics illustrating the procedure to generate the *slo*^E366G^ and *slo*^*loxP*^ alleles via ends-out homologous recombination (A) and method for molecular validation (B). The *slo* locus is shown. Grey boxes represent 5’ and 3’ untranslated regions, black boxes represent both constitutive and alternatively spliced coding exons. Region encompassed by targeting arms is denoted by the grey dashed box. Linearized recombinogenic arms (Arm 1 and Arm 2, separated by *loxP* and mini-*white*^+^ sequences) were liberated from a genomic *p[w25*.*2]* insertion as described in Supplemental Ref. 2. Potential recombinants were initially identified by the presence of non-white eye colour due to the mini-*white*^+^ marker, then validated through PCR via the strategy shown in (B). Validated recombinants are denoted by a PCR product of approximately 3.3 kb amplified from single-fly genomic DNA that was absent in non-recombinant control DNA (-ve control).

**Supplemental Fig. 2.**
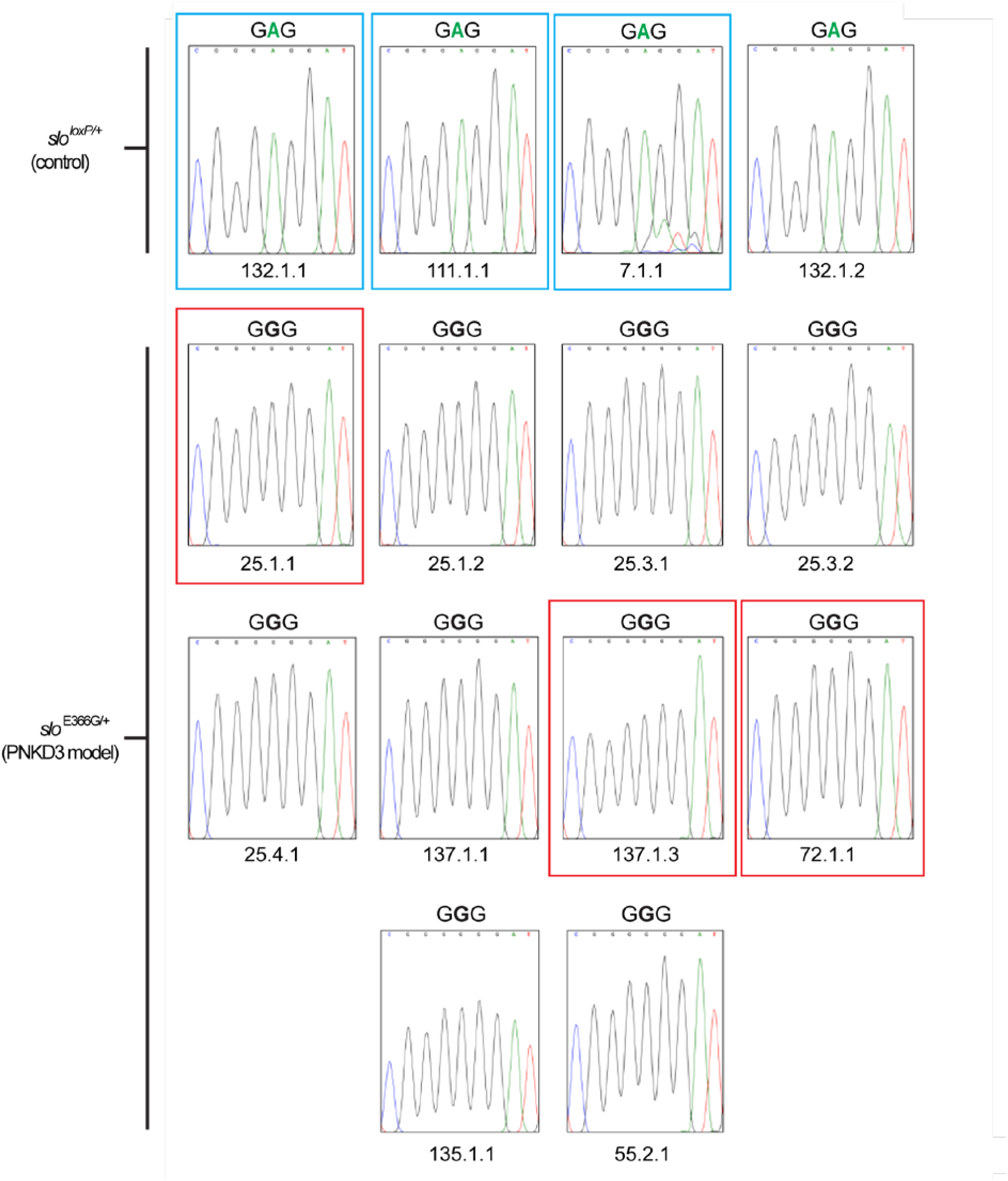
Sequence validation of four independent *slo*^*loxP*^ recombinant lines harbouring the genomically encoded glutamic acid (GAG) at position 366 and ten independent *slo*^E366G^ recombinant lines harbouring the artificially introduced glycine (GGG) at the same position. Lines selected for subsequent out-crossing into the iso31 background are noted in blue and red.

**Supplemental Fig. 3.**
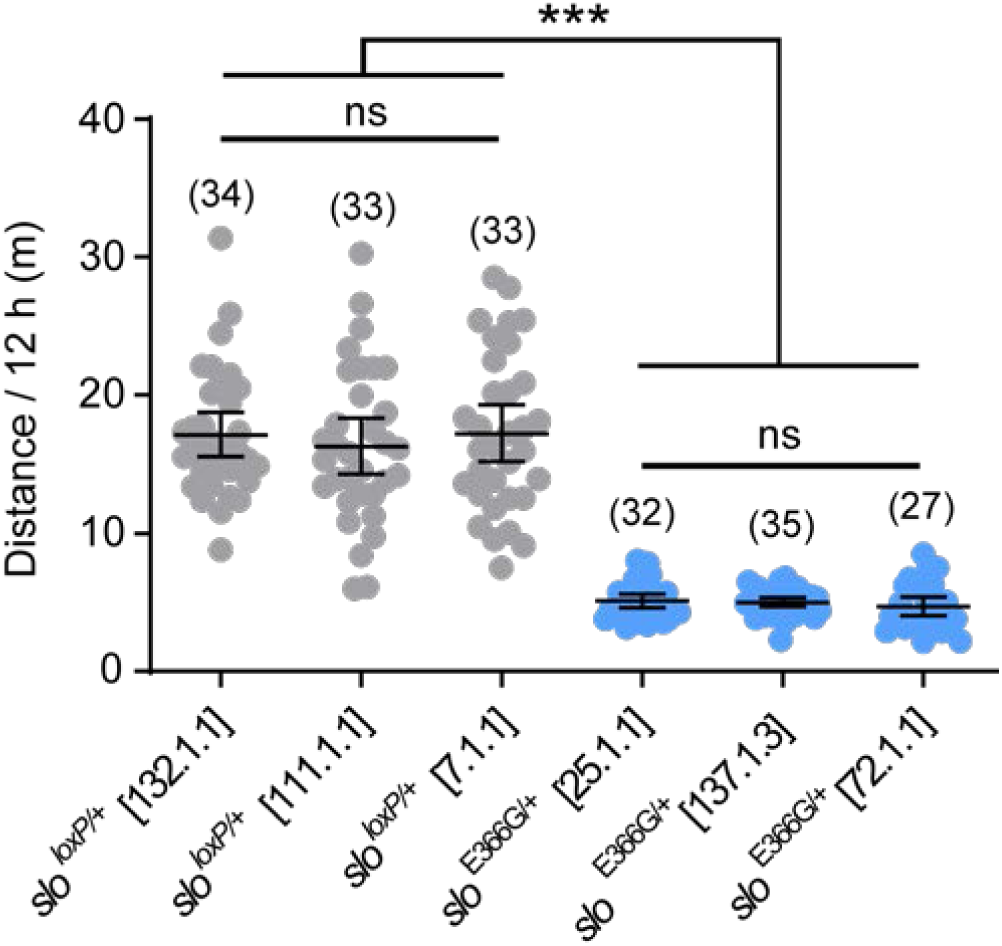
Total distance travelled as measured by the DART system over 12 h in adult males from 3 independent *slo*^*loxP*/+^ and *slo*^E366G/+^ lines. Dots represent individual flies; n-values are noted. Error bars: mean and 95% Confidence Interval (CI). ***p<0.0005, ns – p>0.05, Kruskal-Wallis test with Dunn’s post-hoc test.

**Supplemental Fig. 4.**
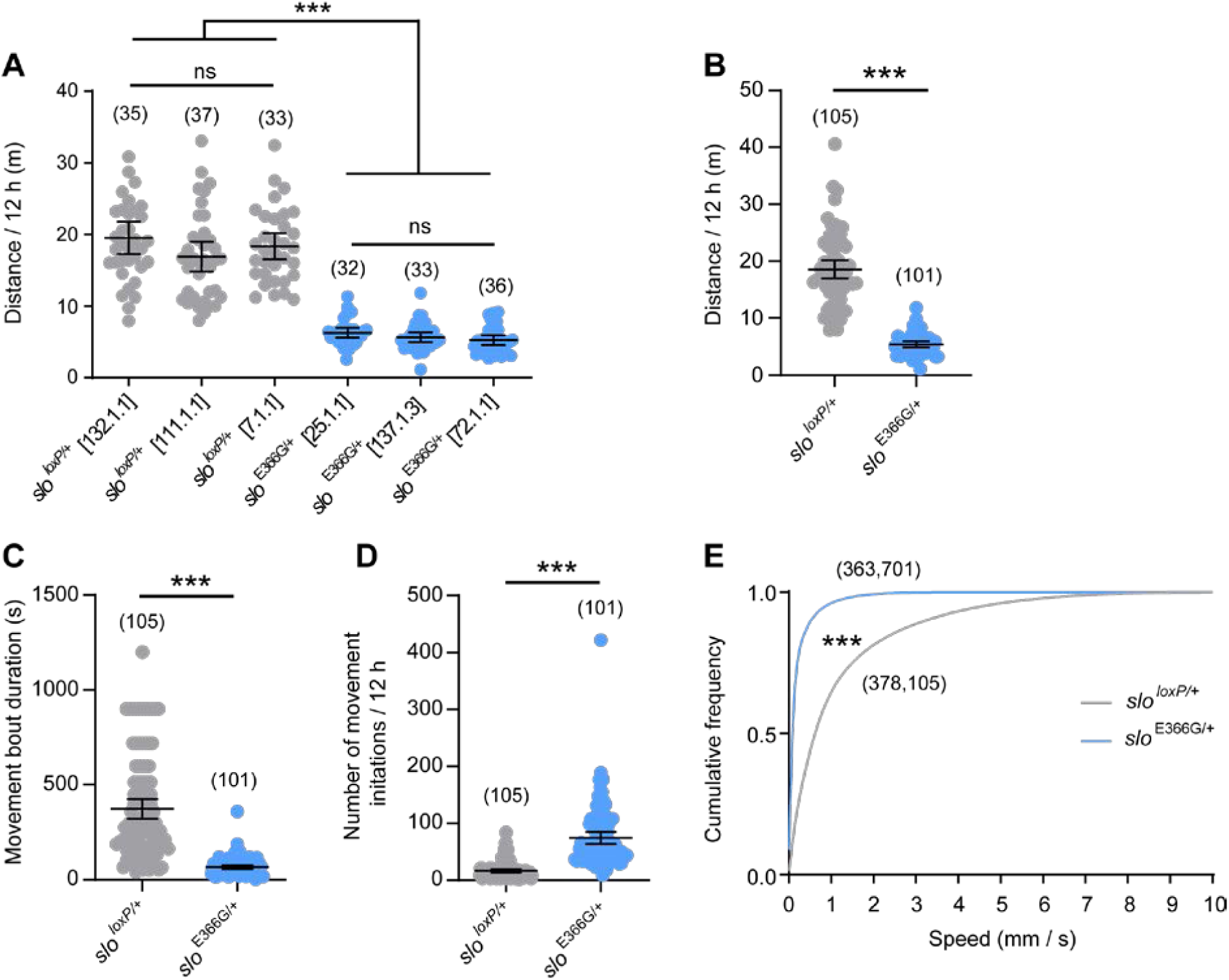
(A-E) Movement parameters in adult female *slo*^*loxP*/+^ and *slo*^E366G/+^ flies measured by the DART system, as described in Fig. 1. Total distance travelled as measured by the DART system over 12 h in adult females from 3 independent *slo*^*loxP*/+^ and *slo*^E366G/+^ lines are shown in (A). (B-E) show pooled data from the 3 *slo*^*loxP*/+^ and *slo*^E366G/+^ lines. Dots represent individual flies; n-values are noted. Error bars: mean and 95% Confidence Interval (CI). ***p<0.0005, ns – p>0.05, Kruskal-Wallis test with Dunn’s post-hoc test (A), Mann-Whitney U-test (B-D) or Kolmogorov-Smirnov test (E).

**Supplemental Fig. 5.**
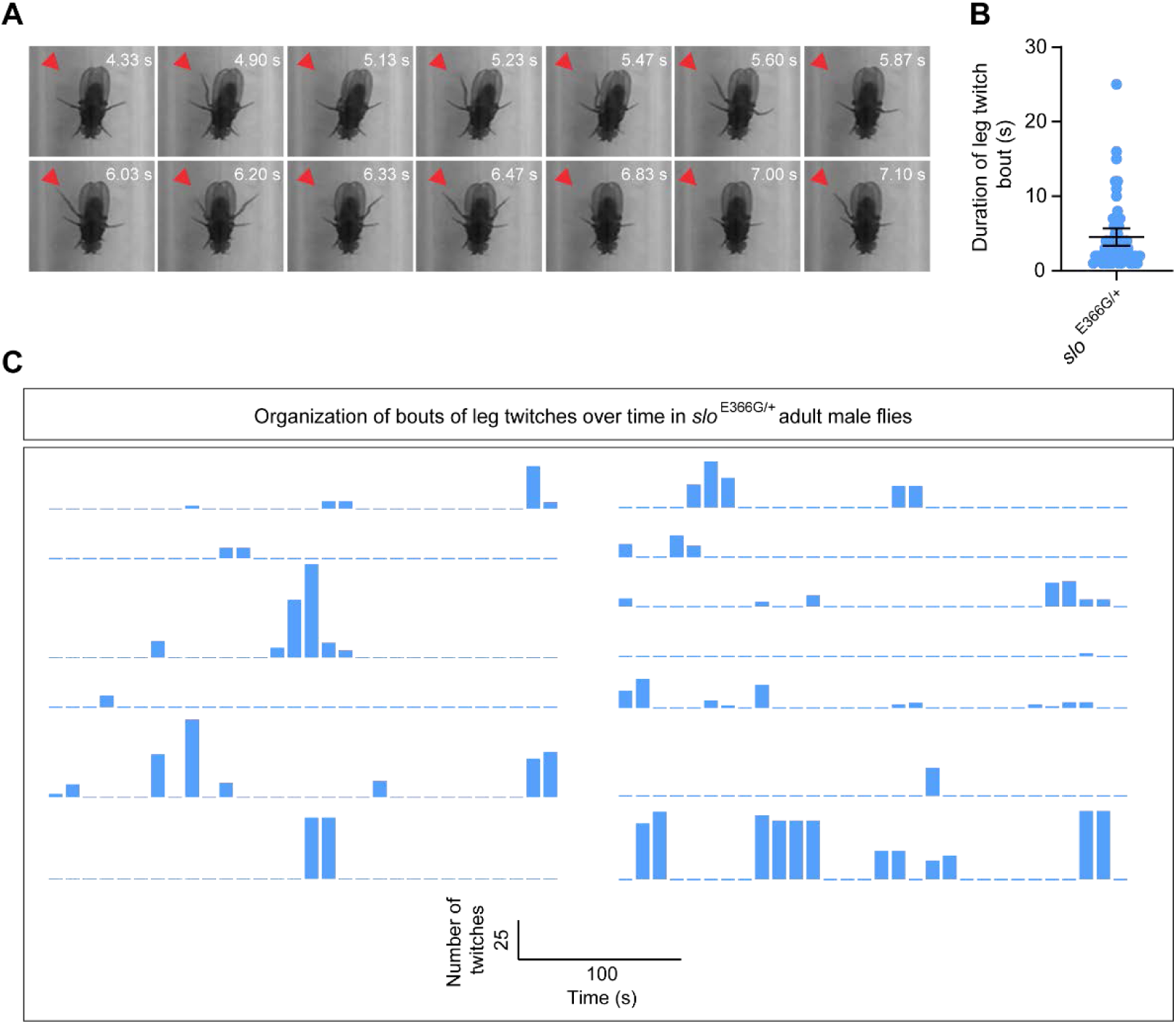
(A) Example still images of a single *slo*^E366G/+^ male exhibiting sporadic leg twitches. Arrow points to rapid extensions and contractions of the rear right leg. (B) Mean duration of individual bouts of leg twitches in *slo*^E366G/+^ males. n = 58 bouts across 13 flies. Error bars, 95% CI. (C) Occurrence of leg twitch bouts over time in *slo*^E366G/+^ males. Each line represents a single in *slo*^E366G/+^ male over a 5 min period, separated into dashed 10 s bins. Where a bout overlapped 10 s bins, the total number of leg twitches was divided evenly between the two bins. Leg twitches were clustered into defined bouts rather than spread uniformly across the 5 min period.

**Supplemental Fig. 6.**
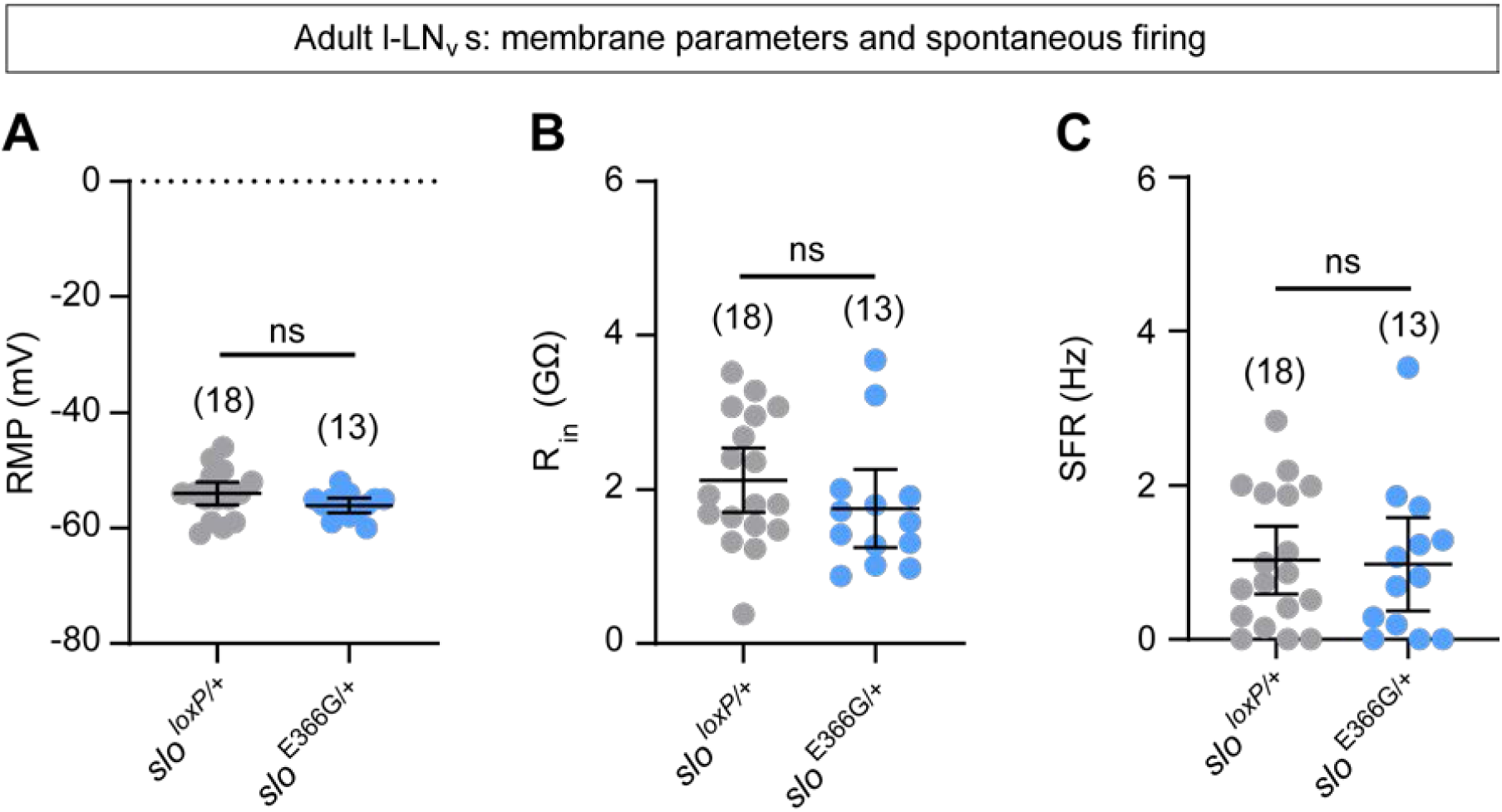
Resting membrane potential (RMP) (A), membrane resistance (R_in_) (B) and spontaneous firing rate (SFR) (C) in adult male *slo*^*loxP*/+^ or *slo*^E366G/+^ l-LN_v_s recorded at ZT18-20. Each dot represents a recording from an individual neuron; n-values are noted. Recordings were obtained from > 9 flies per genotype. Error bars: mean and 95% Confidence Interval (CI). ns – p>0.05, unpaired t-test with Welch’s correction (A) or Mann-Whitney U-test (B, C).

**Supplemental Fig. 7.**
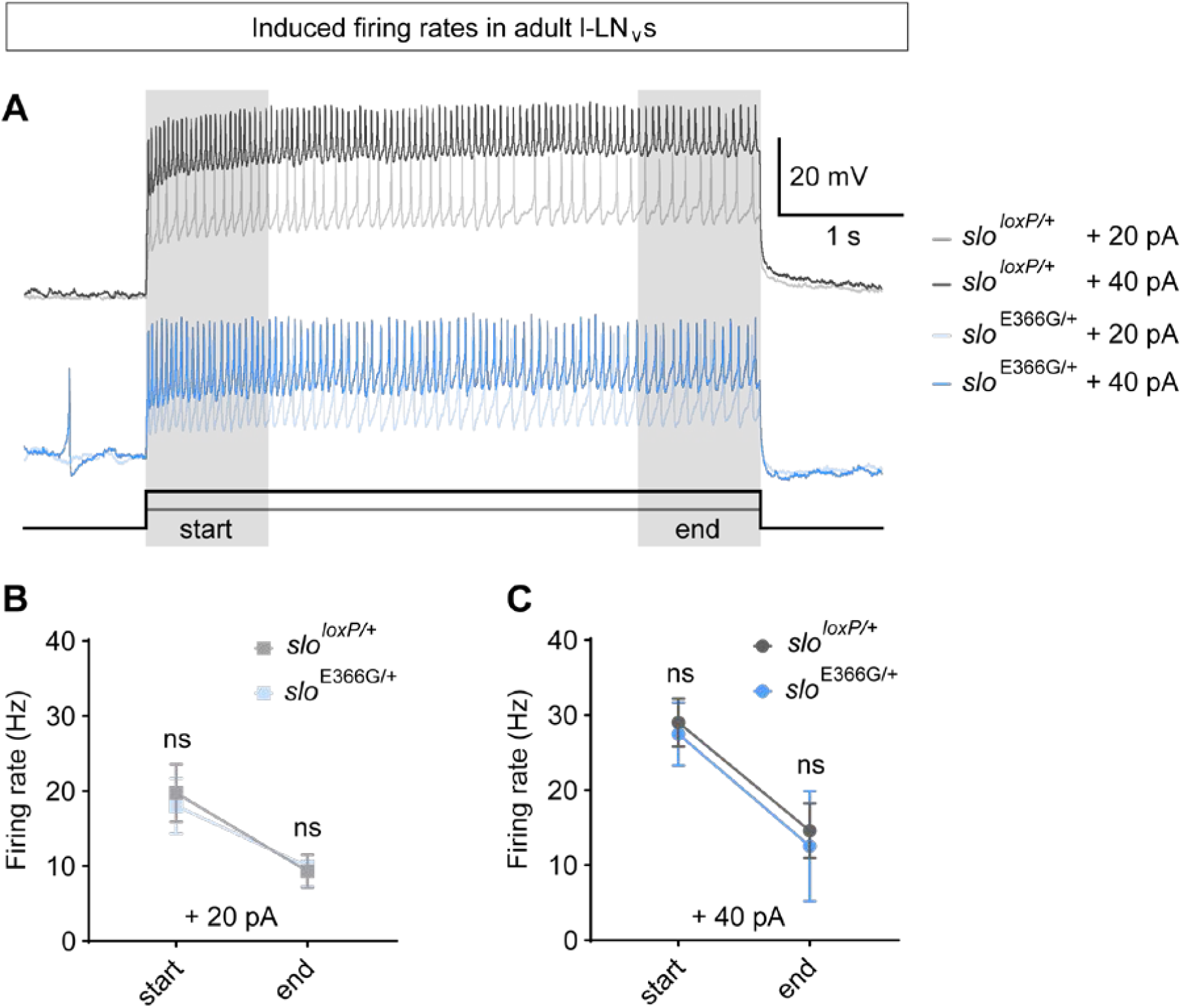
(A) Representative traces showing AP firing in adult male l-LN_v_s from the *slo*^*loxP*/+^ or *slo*^E366G/+^ backgrounds induced by either +20 pA or +40 pA current injections. (B-C) Mean firing rates in adult male l-LN_v_s from the *slo*^*loxP*/+^ or *slo*^E366G/+^ backgrounds at the start or end of AP trains (as indicated in A) induced by +20 pA or +40 pA current injections. N-values are as follows: +20 pA *slo*^*loxP*/+^ - n = 14, +20 pA *slo*^E366G/+^ - n = 17, +40 pA *slo*^*loxP*/+^ - n = 12, +40 pA *slo*^E366G/+^ - n = 15. Error bars: Standard Deviation (SD). ns – p>0.05, Mann-Whitney U-test (+20 pA start, +40 pA start, +40 pA end) or unpaired t-test with Welch’s correction (+20 pA end).

**Supplemental Video 1**. Video shows representative examples of three *slo*^*loxP*/+^ (left) or *slo*^E366G/+^ (right) adult male flies freely moving in glass tubes containing an agar-sucrose food source. Tubes are horizontally placed. Note the more rapid speed and increased continuity of movement of *slo*^*loxP*/+^ compared to *slo*^E366G/+^ males. Subsequent zoom of the left-hand *slo*^E366G/+^ male illustrates a bout of unilateral leg-twitches, similar instances of which were frequently observed in *slo*^E366G/+^ but not *slo*^*loxP*/+^ flies (Fig. 1G).

## Notes

**Financial disclosure:** The authors declare no conflict of interest.

**Funding sources for study:** This study was supported by a Wellcome Strategic Award (WT104033AIA) to DMK and JECJ; a MRC New Investigator Grant (MR/P012256/1) to JECJ; a Wellcome PhD studentship award to PK (109003/Z/15/Z), a MRC Programme Grant to DMK (MR/L01095X/1), and a Leverhulme Trust grant (RPG-2016-318) to JJLH.

## References

1. Erro R, Bhatia KP, Espay AJ, et al. The epileptic and nonepileptic spectrum of paroxysmal dyskinesias: Channelopathies, synaptopathies, and transportopathies. Mov Disord 2017;32(3):310–18. doi: 10.1002/mds.26901 [published Online First: 2017/01/17]

2. Gardiner AR, Jaffer F, Dale RC, et al. The clinical and genetic heterogeneity of paroxysmal dyskinesias. Brain 2015;138(Pt 12):3567–80. doi: 10.1093/brain/awv310 [published Online First: 2015/11/26]

3. Silveira-Moriyama L, Kovac S, Kurian MA, et al. Phenotypes, genotypes, and the management of paroxysmal movement disorders. Dev Med Child Neurol 2018;60(6):559–65. doi: 10.1111/dmcn.13744 [published Online First: 2018/03/31]

4. Bailey CS, Moldenhauer HJ, Park SM, et al. KCNMA1-linked channelopathy. J Gen Physiol 2019;151(10):1173–89. doi: 10.1085/jgp.201912457 [published Online First: 2019/08/21]

5. Du W, Bautista JF, Yang H, et al. Calcium-sensitive potassium channelopathy in human epilepsy and paroxysmal movement disorder. Nat Genet 2005;37(7):733–8. doi: 10.1038/ng1585 [published Online First: 2005/06/07]

6. Griguoli M, Sgritta M, Cherubini E. Presynaptic BK channels control transmitter release: physiological relevance and potential therapeutic implications. J Physiol 2016;594(13):3489–500. doi: 10.1113/JP271841 [published Online First: 2016/03/13]

7. Bean BP. The action potential in mammalian central neurons. Nat Rev Neurosci 2007;8(6):451–65. doi: 10.1038/nrn2148 [published Online First: 2007/05/22]

8. Kadas D, Ryglewski S, Duch C. Transient BK outward current enhances motoneurone firing rates during Drosophila larval locomotion. J Physiol 2015;593(22):4871–88. doi: 10.1113/JP271323 [published Online First: 2015/09/04]

9. Ford KJ, Davis GW. Archaerhodopsin voltage imaging: synaptic calcium and BK channels stabilize action potential repolarization at the Drosophila neuromuscular junction. J Neurosci 2014;34(44):14517–25. doi: 10.1523/JNEUROSCI.2203-14.2014 [published Online First: 2014/10/31]

10. Diez-Sampedro A, Silverman WR, Bautista JF, et al. Mechanism of increased open probability by a mutation of the BK channel. J Neurophysiol 2006;96(3):1507–16. doi: 10.1152/jn.00461.2006 [published Online First: 2006/06/02]

11. Yang J, Krishnamoorthy G, Saxena A, et al. An epilepsy/dyskinesia-associated mutation enhances BK channel activation by potentiating Ca2+ sensing. Neuron 2010;66(6):871–83. doi: 10.1016/j.neuron.2010.05.009 [published Online First: 2010/07/14]

12. Zhang ZB, Tian MQ, Gao K, et al. De novo KCNMA1 mutations in children with early-onset paroxysmal dyskinesia and developmental delay. Mov Disord 2015;30(9):1290–2. doi: 10.1002/mds.26216 [published Online First: 2015/07/22]

13. Telford MJ, Budd GE, Philippe H. Phylogenomic Insights into Animal Evolution. Curr Biol 2015;25(19):R876–87. doi: 10.1016/j.cub.2015.07.060 [published Online First: 2015/10/07]

14. Rong YS, Titen SW, Xie HB, et al. Targeted mutagenesis by homologous recombination in D. melanogaster. Genes Dev 2002;16(12):1568–81. doi: 10.1101/gad.986602 [published Online First: 2002/06/25]

15. Faville R, Kottler B, Goodhill GJ, et al. How deeply does your mutant sleep? Probing arousal to better understand sleep defects in Drosophila. Sci Rep 2015;5:8454. doi: 10.1038/srep08454 [published Online First: 2015/02/14]

16. Koh K, Joiner WJ, Wu MN, et al. Identification of SLEEPLESS, a sleep-promoting factor. Science 2008;321(5887):372–6. doi: 10.1126/science.1155942 [published Online First: 2008/07/19]

17. Tabuchi M, Monaco JD, Duan G, et al. Clock-Generated Temporal Codes Determine Synaptic Plasticity to Control Sleep. Cell 2018;175(5):1213–27 e18. doi: 10.1016/j.cell.2018.09.016 [published Online First: 2018/10/16]

18. Jepson JE, Shahidullah M, Lamaze A, et al. dyschronic, a Drosophila homolog of a deaf-blindness gene, regulates circadian output and Slowpoke channels. PLoS Genet 2012;8(4):e1002671. doi: 10.1371/journal.pgen.1002671 [published Online First: 2012/04/26]

19. Mburu P, Mustapha M, Varela A, et al. Defects in whirlin, a PDZ domain molecule involved in stereocilia elongation, cause deafness in the whirler mouse and families with DFNB31. Nat Genet 2003;34(4):421–8. doi: 10.1038/ng1208 [published Online First: 2003/07/02]

20. Pfeiffenberger C, Lear BC, Keegan KP, et al. Locomotor activity level monitoring using the Drosophila Activity Monitoring (DAM) System. Cold Spring Harb Protoc 2010;2010(11):pdb prot5518. doi: 10.1101/pdb.prot5518 [published Online First: 2010/11/03]

21. Lee US, Cui J. {beta} subunit-specific modulations of BK channel function by a mutation associated with epilepsy and dyskinesia. J Physiol 2009;587(Pt 7):1481–98. doi: 10.1113/jphysiol.2009.169243 [published Online First: 2009/02/11]

22. Wang B, Rothberg BS, Brenner R. Mechanism of increased BK channel activation from a channel mutation that causes epilepsy. J Gen Physiol 2009;133(3):283–94. doi: 10.1085/jgp.200810141 [published Online First: 2009/02/11]

23. Brenner R, Chen QH, Vilaythong A, et al. BK channel beta4 subunit reduces dentate gyrus excitability and protects against temporal lobe seizures. Nat Neurosci 2005;8(12):1752–9. doi: 10.1038/nn1573 [published Online First: 2005/11/02]

24. Montgomery JR, Meredith AL. Genetic activation of BK currents in vivo generates bidirectional effects on neuronal excitability. Proc Natl Acad Sci U S A 2012;109(46):18997–9002. doi: 10.1073/pnas.1205573109 [published Online First: 2012/11/01]

25. Raffaelli G, Saviane C, Mohajerani MH, et al. BK potassium channels control transmitter release at CA3-CA3 synapses in the rat hippocampus. J Physiol 2004;557(Pt 1):147–57. doi: 10.1113/jphysiol.2004.062661 [published Online First: 2004/03/23]

26. Renn SC, Park JH, Rosbash M, et al. A pdf neuropeptide gene mutation and ablation of PDF neurons each cause severe abnormalities of behavioral circadian rhythms in Drosophila. Cell 1999;99(7):791–802. doi: 10.1016/s0092-8674(00)81676-1 [published Online First: 2000/01/05]

27. Ruben M, Drapeau MD, Mizrak D, et al. A mechanism for circadian control of pacemaker neuron excitability. J Biol Rhythms 2012;27(5):353–64. doi: 10.1177/0748730412455918 [published Online First: 2012/09/27]

## Supplemental References

1. Rong YS, Titen SW, Xie HB, et al. Targeted mutagenesis by homologous recombination in D. melanogaster. Genes Dev 2002;16(12):1568–81. doi: 10.1101/gad.986602 [published Online First: 2002/06/25]

2. Staber CJ, Gell S, Jepson JE, et al. Perturbing A-to-I RNA editing using genetics and homologous recombination. Methods Mol Biol 2011;718:41–73. doi: 10.1007/978-1-61779-018-8_3 [published Online First: 2011/03/04]

3. Langley CH, Stevens K, Cardeno C, et al. Genomic variation in natural populations of Drosophila melanogaster. Genetics 2012;192(2):533–98. doi: 10.1534/genetics.112.142018 [published Online First: 2012/06/08]

4. Pool JE, Corbett-Detig RB, Sugino RP, et al. Population Genomics of sub-saharan Drosophila melanogaster: African diversity and non-African admixture. PLoS Genet 2012;8(12):e1003080. doi: 10.1371/journal.pgen.1003080 [published Online First: 2013/01/04]

5. Chen KF, Lowe S, Lamaze A, et al. Neurocalcin regulates nighttime sleep and arousal in Drosophila. Elife 2019;8 doi: 10.7554/eLife.38114 [published Online First: 2019/03/14]

6. Buhl E, Bradlaugh A, Ogueta M, et al. Quasimodo mediates daily and acute light effects on Drosophila clock neuron excitability. Proc Natl Acad Sci U S A 2016;113(47):13486–91. doi: 10.1073/pnas.1606547113 [published Online First: 2016/11/09]

7. Ruben M, Drapeau MD, Mizrak D, et al. A mechanism for circadian control of pacemaker neuron excitability. J Biol Rhythms 2012;27(5):353–64. doi: 10.1177/0748730412455918 [published Online First: 2012/09/27]

8. Tabuchi M, Monaco JD, Duan G, et al. Clock-Generated Temporal Codes Determine Synaptic Plasticity to Control Sleep. Cell 2018;175(5):1213–27 e18. doi: 10.1016/j.cell.2018.09.016 [published Online First: 2018/10/16]

9. Telford MJ, Budd GE, Philippe H. Phylogenomic Insights into Animal Evolution. Curr Biol 2015;25(19):R876–87. doi: 10.1016/j.cub.2015.07.060 [published Online First: 2015/10/07]

